# Multi-species polyploidization, chromosome shuffling, and genome extraction in *Zea/Tripsacum* hybrids

**DOI:** 10.1101/2023.02.02.526764

**Authors:** Muhammad Zafar Iqbal, Xiaodong Wen, Lulu Xu, Yanli Zhao, Jing Li, Weiming Jiang, Mingjun Cheng, Huaxiong Li, Yinzheng Li, Xiaofeng Li, Ruyu He, Jianmei He, Asif Ali, Yan Peng, Tingzhao Rong, Qilin Tang

## Abstract

By hybridization and special sexual reproduction, we sequentially aggregated *Zea mays, Zea perennis,* and *Tripsacum dactyloides* in an allohexaploid, backcrossed it with maize, derived self-fertile allotetraploids of maize and *Z. perennis* by natural genome extraction, extended their first six selfed generations, and finally constructed amphitetraploid maize using nascent allotetraploids as a genetic bridge. Transgenerational chromosome inheritance, subgenomes stability, chromosome pairings and rearrangements, and their impacts on an organism’s fitness were investigated by fertility phenotyping and molecular cytogenetics techniques GISH and FISH. Results showed that diversified sexual reproductive methods produced highly differentiated progenies (2n=35-84) with varying proportions of subgenomic chromosomes, of which one individual (2n=54, MMMPT) overcame self-incompatibility barriers and produced self-fertile nascent near-allotetraploid by preferentially eliminating *Tripsacum* chromosomes. Nascent near-allotetraploid progenies showed persistent chromosome changes, intergenomic translocations, and rDNA variations for at least up to the first six selfed generations; however, ploidy tended to stabilize at near-tetraploid level (2*n*=40) with full integrity of 45SrDNA pairs, and a trend of decreasing variations by advancing generations with an average of 25.53, 14.14, and 0.37 maize, *Z. perennis,* and *T. dactyloides* chromosomes, respectively. The mechanisms for three genome stabilities and karyotype evolution for formatting new polyploid species were discussed.

## Introduction

According to the classical view, any ancestral species in the tree of life may divide and separate its gene pool into different populations, and then these populations differentiate into different species (Dobzhansky 1937; Grant 1981). The creation of new species is certainly not conscious or purposeful behavior of old species, and the genome of any descendant individual is one of the random combinations separated from the gene pool of the ancestral separated population. The diversity of the gene pool of the divided population determines the diversity of the individual genetic combination of the descendant; natural selection selects only favorable individuals among various randomly separated combinations and interacts with the environment screening to form a variety of new species. Based on the evolutionary tree, the evolution of species leads to the diversity of species, but for subsequent offspring species, the division of ancestral genetic material leads to the relative reduction of genetic diversity of individual offspring species (Chen et al. 2021; Yu and Li 2022). Furthermore, the irreversible law of biological evolution describes that certain types of evolved organisms cannot restore their ancestry, and the extinct type cannot reappear (Louis Dollo, 1893).

However, hybridization and the special aspects of sexual reproduction may have contributed to species diversification and crop improvements (Abbott et al. 2013; De Storme and Mason 2014; Rieseberg and Willis 2007). When special sexual reproduction and hybridization are used to purposefully bring different species or genera together for genetic recombinations and break the species and genus boundaries, they expand genetic variations so that realizing the breeding significance of improving old species or may even create new species by reversing the species to their primitive forms, reproduce extinct types, or through creating new unique combinations (Alix et al. 2017; Mason 2017; Mason and Batley 2015; Udall and Wendel 2006). Notably, some scientists have recently proposed to domesticate wild relatives of crops for elite traits (Fernie and Yan 2019; Yu et al. 2021; Zsögön et al. 2018). Polyploids generally show higher genome heterozygosity and allelic diversity with better physiological adoptability than diploids, thus creating polyploids and their genetic breeding has long been one of the important means to increase genetic variations and trait improvement (Mason and Batley 2015; Udall and Wendel 2006; Van de Peer et al. 2021; Van de Peer et al. 2017).

Maize (*Zea mays* ssp. *mays*, 2n=20) is one of the world’s three most important cereals and is the only cultivated crop among its close relatives of teosinte and *Tripsacum*. All species of the genus *Zea* except *Zea mays* ssp. *mays* are collectively known as teosintes, considered the most probable progenitors of domesticated maize (Ross-Ibarra et al. 2009; Welker et al. 2020). Maize originated in southern Mexico and was first domesticated from teosinte over 6,000 years ago (Da Fonseca et al. 2015). *Zea perennis* (PPPP; 2n=40) is the only tetraploid teosinte and is considered the most primitive species of the genus. It possesses many attractive breeding traits such as perenniality, cold, and flood tolerance. Species of the genus *Tripsacum* are more distantly related to maize and teosinte, and the *Tripsacum* is considered the sister genera of the *Zea. Tripsacum* and *Zea* diverged from a common ancestor fewer than 1.2 million years ago (Ross-Ibarra et al. 2009; Welker et al. 2020) and have shared a common polyploidy event (Estep et al. 2014). *Tripsacum* owns excellent genes for biotic and abiotic stress tolerance and apomixis – lacking in maize, thus have often been hybridized with maize (Harlan and Wet 1977; Kindiger and Dewald 1996; Mangelsdorf and Reeves 1931; Yan et al. 2020). Natural gene flow from *Z. perennis* or *Tripsacum* into cultivated maize is seldom, if any. However, artificial hybridization is possible (Harlan and De Wet 1977; Shaver 1964; Tang et al. 2005; Yan et al. 2020).

Maize tetraploid hybrids show elevated vigor and progressive heterosis that diploid maize hybrids lack (Riddle and Birchler 2008; Washburn et al. 2019). Several maize autopolyploids, “from monoploid to octoploid”, have been synthesized (Auger et al. 2004; Birchler and Veitia 2012; Chase 1980; Kato and Birchler 2006; Rhoades and Dempsey 1966; Riddle et al. 2006; Yao et al. 2013; Yao et al. 2011) and the result showed that strictly homozygous material imposed adverse effects on the fertility and viability of polyploid individuals and decline in stature by increasing ploidy. Maximizing heterozygosity either by restructuring the genomes of autopolyploid or allopolyploidization improved bivalent pairing and increased plant stature (Doyle 1986; Rhoades and Dempsey 1966; Riddle and Birchler 2008; Shaver 1963). However, this gain is challenging to utilize for agronomic purposes due to the quadrivalent pairing of chromosomes and double reduction. These barriers can be theoretically overcome by inducing apomixes or chromosomal-level diploidization, but so far, all in vivo attempts have achieved minimal success.

Species evolution is the separation and formation of various branch-related species under natural conditions. If we use hybridization and special sexual reproduction methods to aggregate the genomes of key close relatives of phylogenetic trees and then do genome shuffling, can it create new species that did not appear before? If so, the formation of new species or the expansion of plant adaptability can be completed almost immediately. This celerity can be achieved through hybridization and special reproductive methods that aggregate genomes of different existing species rather than through long-term evolutionary changes in DNA sequences. This study used hybridization and 2n+n special sexual reproduction to construct a tri-species intergeneric hybrid (MMPPTT, 2n=74) among maize, *Z. perennis*, and *T. dactyloides* and then crossed it with maize to synthesize a series of allopolyploids among three or two parental species by sequential processes of backcrossing and self-breeding. Thus, using hybridization and special sexual reproduction to aggregate the genomes of three close relatives, we restore and reproduce genomes similar to ancestral types. Moreover, it provides a new approach to crop breeding by creating rare materials or obtaining useful genes by aggregating multiple close relatives.

## Material and Methods

### Plant materials

The plant material includes a tri-genomic allohexaploid hybrid (2n=74, MMTTPP) formed by hybridizing (*Zea mays* ssp. *mays*, 2n=4x=40 x *Tripsacum dactyloides*, 2n=4x=72) x *Zea perennis*, 2n=4x=40, which on backcrossing to diploid maize (2n=2x=20) produced a series of allopolyploid progenies (2n=35-84) harboring three parental species genomes in different proportions, of which one individual overcame a self-incompatibility barrier, thereby was used to generate its the first six-selfed generations by selecting and propagating the most fertile individuals for the next generation. Finally, it was hybridized to two autotetraploid maize inbreds as a genetic bridge to construct the genomes of tetraploid maize inbreds for improved fertility. Two independent populations of the genome-constructed tetraploid maize (amphitetraploid maize) were extended up to the first three selfed generations (See Suppl. Material and Methods for detail)

### Pollen viability and seed setting percentage

Pollen viability was assessed by the I_2_-KI staining method, and seed setting percentage was calculated by the total number of seeds per cob/total number of glumes per cob multiplied by 100. (See detail in Method S1).

### Chromosome counts, mitotic and meiotic GISH, FISH, and chromosomes karyotyping

Carbol fuchsin staining was used to count the total chromosomes number/cell. Genomic in Situ Hybridization (GISH) was used to discriminate mitotic and meiotic chromosomes of different species within a single nucleus and to identify inter-genomic chromosome rearrangements. Fluorescent in situ Hybridization (FISH) with Cent. C, 45SrDNA, and 5S genes were employed to track the evolution of ribosomal DNA and chromosome rearrangements. Individual chromosomes were identified by karyotyping using DRAWID and Image Pro Plus software. Photoshop was used to process chromosome pictures and adjust signal strength. The detailed procedures are described in Method S1.

### Analysis of ITS cloned sequences

The ITS (ITS1-5.8S–ITS2) region of rDNA of nascent near-allotetraploids and parents were PCR amplified, cloned, and sequenced. The detailed procedures are given in Method S1.

### Amplified fragment length polymorphism (AFLP)

AFLP markers are used to detect genetic transfer from *T. dactyloides* into nascent near-allotetraploids along with nature and time span of genomic changes. Operational detail is provided in Method S1.

### Statistical analysis

Statistics were performed by the data analysis function of Microsoft Excel, IBM SPSS21, and PAST. Details of statistical analysis are given in Method S1.

### Data availability

The authors confirm that the major data supporting the findings of this study are available within the article and its online supplementary file (https://figshare.com/s/3d7ea36fa21e7240ad4c). Raw data are available from the corresponding author on request.

## Results

### Diversified sexual reproductive methods produced highly diversified germplasm

This research group had already synthesized an intergeneric tri-species hybrid “MTP” by hybridizing (tetraploid maize x tetraploid *T. dactyloides*) x tetraploid *Z. perennis* (Yan et al. 2020). It was formed by fertilization of an unreduced gamete of the F1 hybrid with a reduced gamete of *Z. perennis* and harbored chromosome complements 2n=74=20M+20P+34T (M, P, and T stand for maize, *Z. perennis,* and *T. dactyloides*). Karyotype analysis showed that MTP had complete diploid sets of maize and *Z. perennis* chromosomes and a diploid set of *T. dactyloides* chromosomes without chromosomes 32 and 35 (**Figure S2a**). By backcrossing of MTP (2n=74) to diploid maize (2n=20), 47 chromosomes were theoretically expected; despite this, chromosomes ranged from 2n=35-84 in the population of the first backcross generation (BC1) (**Figure S3**). GISH analysis revealed that 5 sexual reproduction pathways were involved to produce these highly diversified populations (**Figure S4**). 1) Parthenogenesis (n): reduced gametes of MTP directly developed into seeds without fertilization, harboring 35-38 chromosomes per plant, and M, P and T chromosomes ranged from 10-12, 8-9, 16-17, respectively (**Figure 1a**). 2) Sexual reproduction (n+n): the population developed by fertilization of reduced gametes from both parents, thus harboring chromosome complements ranging from 2n=44-46 (18-22M+9-11P+15-16T) (**Figure 1b-e**). 3) Polyspermy (n+n+n): the progeny produced by fertilization of a reduced gamete of MTP with two reduced gametes of maize harboring somatic chromosomes complements ranging from 2n=54-57 (28-30M+11P+15-16T) (**Figure 1f-g**). 4) Apomictic mode (2n): the seeds developed directly from unreduced gametes of MTP by omitting fertilization, thereby chromosome complements in this group ranged from 2n=70-74 (19-20M+19-20P+32-34T) (**Figure 1h**). 5) Fertilization of unreduced gametes (2n+n): developed by fertilization of an unreduced gamete of MTP with reduced maize gamete, thus harboring chromosome complement of 2n=84 (30M+20P+34T) (**Figure 1i**). Of 14 BC_1_ individuals, 9 plants harbored translocated chromosomes (range 1 to 4) between maize and *Z. perennis* chromosomes (MP), 2 plants harbored 2 translocations between maize and *Tripsacum*, and two plants had 2 translocations between *Tripsacum* and *Z. perennis* chromosomes (**Figure S4**). Karyotype analysis showed BC1-26 (2n=74) developed by an apomictic pathway harboring the same subgenomic chromosomes as were observed in MTP (**Figure S2a**); BC1-6 (2n=84) formed by fertilization of an unreduced gamete with reduced maize gametes also having the same *T. dactyloides* and *Z. perennis* chromosomes as were observed in MTP, but all maize chromosome had three copies (**Figure S2b)**.

**Figure 1.**
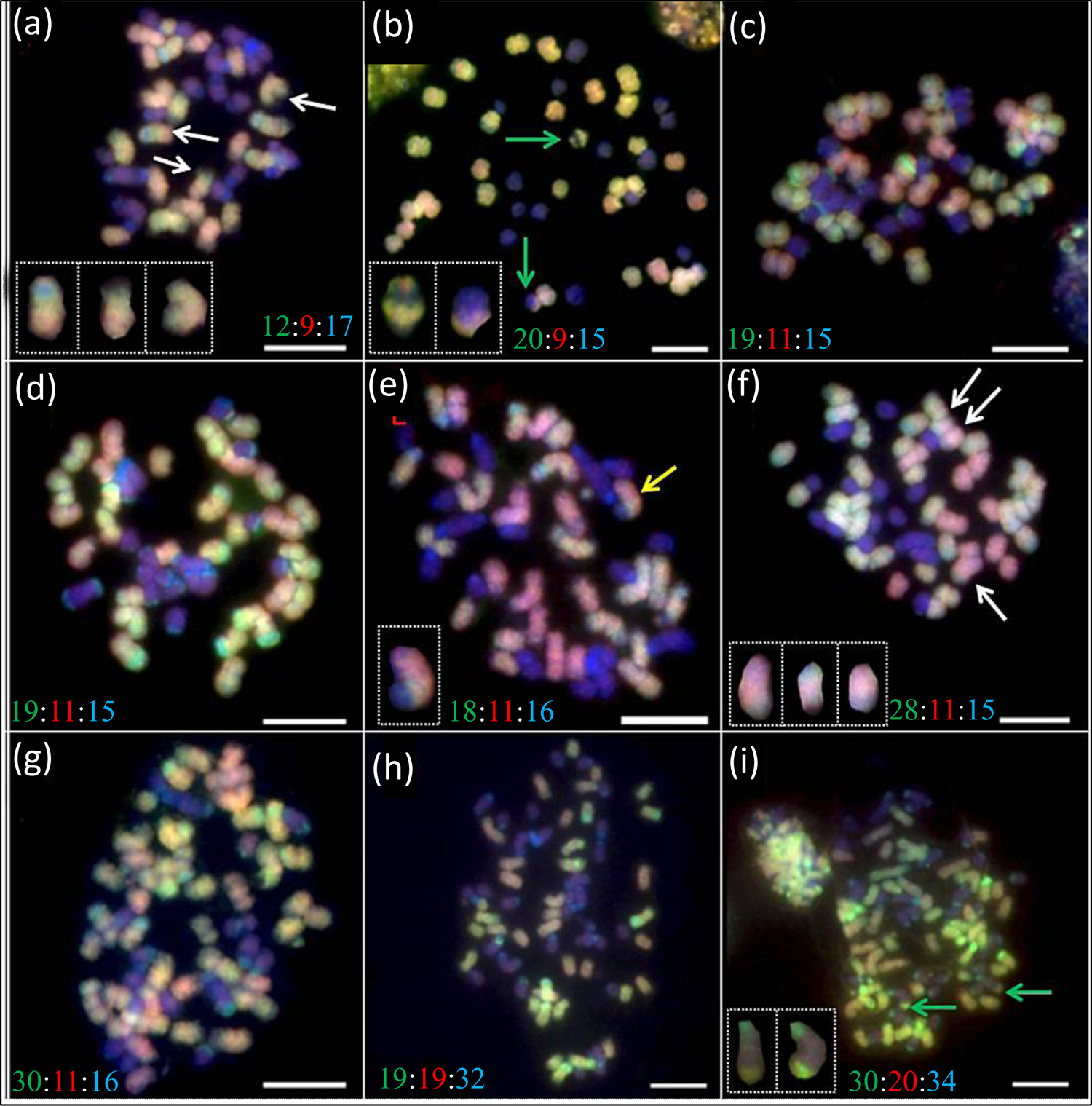
Multi-color GISH analysis of root-tip mitotic cells of the first backcross generation (BC1) of tri-species intergeneric hybrid. Green, red, and blue colors show the chromosomes of maize (Mz), *Z. perennis* (Zp), and T. dactyloides (Tr), respectively. Arrows indicate chromosomal translocations, and rectangular shapes display translated chromosomes within one cell. On insets, green, red, and blue numbers represent chromosomes from Mz, Zp, and Tr genomes, respectively. The bar is 10 um. **(a)**; BC_1_-32. **(b)**; BC_1_-5. **(c)**; BC_1_-19. **(d)**; BC_1_-25. **(e)**; BC_1_-40. **(f)**; BC_1_-31. **(g)**; BC_1_-10. **(h)**; BC_1_-16. **(i)**; BC_1_-6.

The individual developed by other pathways (polyspermy, sexual, and parthenogenesis) having irregular chromosome copy number variations relative to additive parental theoretical values (**Figure S2c and Figure S4**). The polyspermy pathway produced somehow self-fertile progeny with pollen fertility ranging from 7.38% to 37.5 %, while the remaining populations were pollen sterile (**Table S1**). Phenotypically, plants were intermediate between parents (profound tillers, secondary branches, shorter ears) or trended to maize (fewer tillers or without tillers); cobs were shorter with few seed rows, and tillers ranged from 2-48 (Average; 11.7) per plant (**Figure 2a-h and Table S1**). The plants developed by 2n and 2n+n pathways were perennial, showing luxuriant growth; n+n+n trended to maize and n+n were mixed swarms. The population developed by parthenogenesis (*n*) was very weak, showing intermediate phenotypes of three parents, and most individuals could not complete the life cycle. An individual “BC1-31” developed by the n+n+n pathway showed the highest pollen fertility (37.50%), harboring chromosome complements 2n=54 (28M+11P+15T) and had a maize-like plant stature.

**Figure 2.**
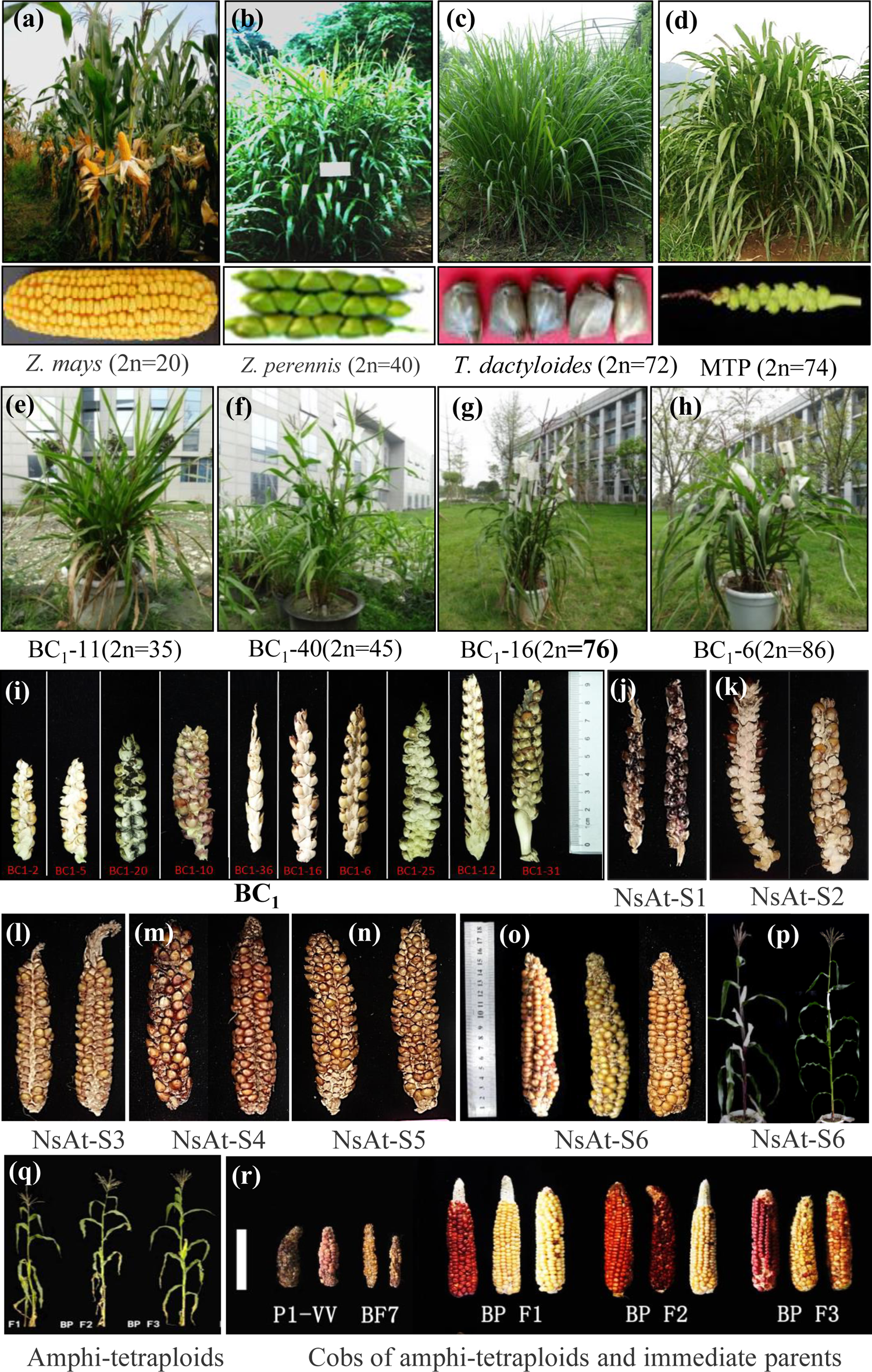
Phenotypic manifestation of parents and sequentially derived nascent near-allotetraploids (NsAt) by subgenomes extraction. **(a-c);** Parental species plants with cobs. **(d);** Tri-species intergeneric allohexaploid hybrid (2*n*=74, MTP) with cob. **(e-h);** Representative backcrossed (MTP x diploid maize) progenies formed by parthenogenesis (n), sexual reproduction (n+n), apomixes (2n), and fertilization of an unreduced gamete, respectively. **(i)** ear morphology of BC1 individuals, (**j-o**) ear morphologies of nascent near-allotetraploids of the first six selfed generations (S1 to S6), respectively, formed by natural genome extraction; **(p)** two representative plants of the two different lines of the six^th^ selfed generation. **(q);** Representative plants of the first three selfed generations (F1-F3) of amphitetraploid maize constructed using nascent near-allotetraploid as genetic bridge materials. **(r)** representative pictures of the cobs of the first three selfed generations (F1-F3) with cobs of immediate parents (bar 10 cm).

### Formation of near-allotetraploid lines by natural genome extraction and chromosome shuffling

To answer a question, what would be the fate of BC1-31 (2n=54=28M+11P+15T), showing 37.50% pollen fertility, if propagated for the next generation? It could either establish new stable fertile allopolyploid lines or become extinct, or if it produces fertile progenies, how do three subgenomes integrate into one nucleus? Consequently, we selfed the BC1-31 and obtained 24 seeds from many pollinated florets. Of them, 16 seeds germinated on planting, and chromosomes of only 10 plants could be identified. Results showed the selfed progeny of BC1-31 (S1) with drastically reduced chromosome numbers ranging from 2n=31-44, with 9 plants of 2n=41-44 (all self-fertile) and 1 plant of 2n=33 (pollen sterile). mc-GISH analysis revealed that BC1 individuals harbored maize M, *Z. perennis*, and *T.dactyloides* chromosomes in the range of 23-28, 5-17, and 1-5 (Mean 25.20M+13.3P+2.50T), respectively. A drastic loss occurred for *Tripsacum* chromosomes, while the loss of some maize chromosomes was compensated by extra copies of *Z. perennis* chromosomes.

Next, we selected the five most fertile plants and propagated the first six selfed generations by selecting the most fertile individuals for every next generation. The populations of the first six selfed generations were self-fertile and did not show any remarkable fecundity at least up to the 6^th^ selfed generation (S1-S6), with a trend to a maize-like plant stature, synchronization in flowering phenology, and an increase in seed rows/cob and cob length with advancing selfed generations (**Figure 2i-o, see detailed graphical representation in Figure S5**). A trend of gradually increasing seed number was observed by advancing the generations, while average seed set and pollen viability initially increased up to the first three selfed generations and then decreased but remained higher or at least similar to that of the S1 population (**Figure 3a-c and Table S2**); however a significant variation existed for pollen viability, seed set, and seed number per plant for individual values within the generations and among generations (**Figure 3a1-c1)**. Branched/apical dominance and paired/single mature spikelet are two major traits differentiating teosinte and maize. Plants architecture of these allopolyploids was more likely similar to maize (**Figure 3e**), showing apical dominance (**Figure 2o**), and cobs had paired mature spikelets (**Figure 2j-n**).

**Figure 3.**
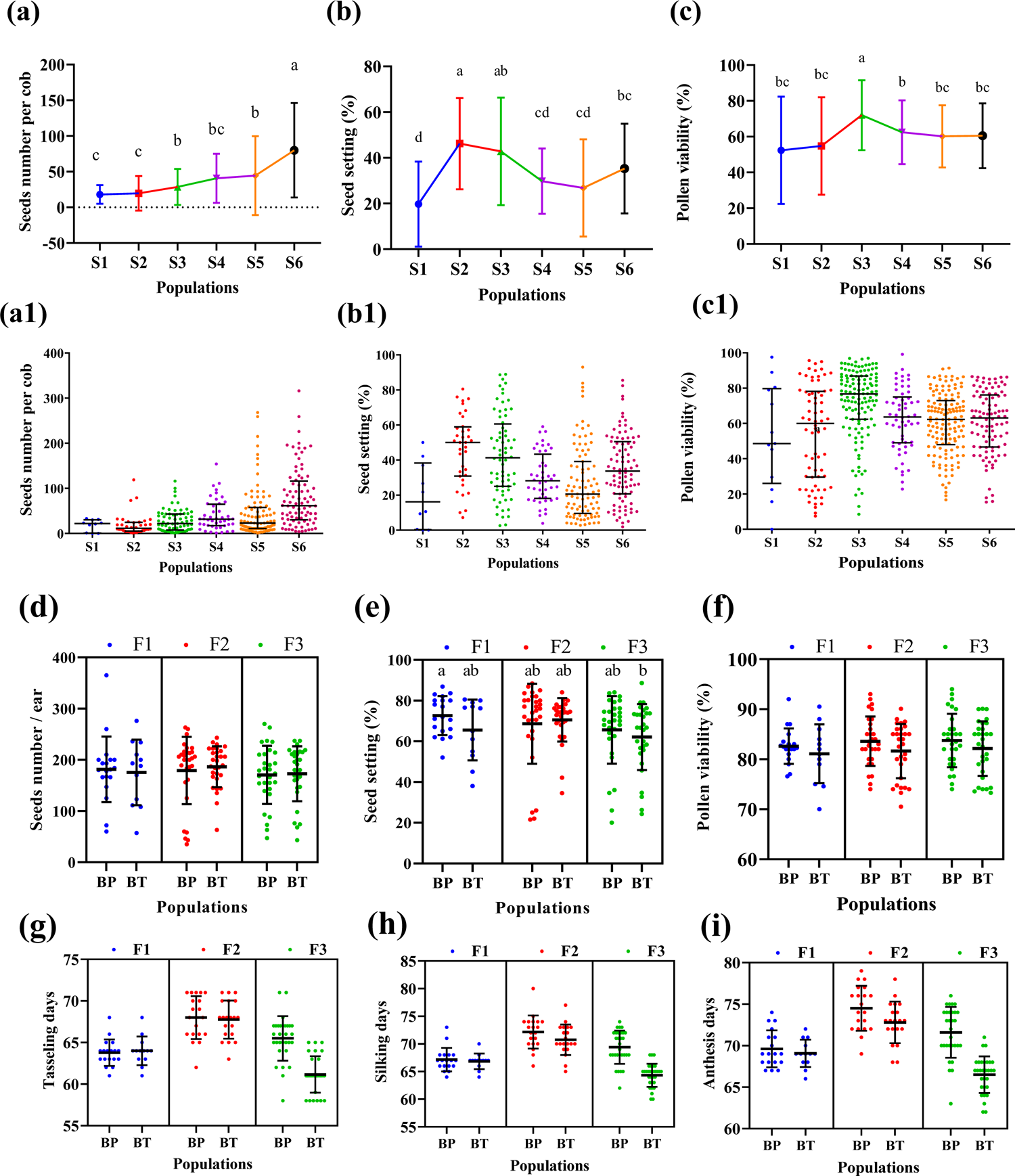
Fertility phenotyping and flowering phenology of derived allotetraploids between maize and *Z. perennis*. **(a-a1), (b-b1),** and **(c-c1)** show the comparison of seed number, seed setting, and pollen viability among six consecutive selfed generations of nascent near-allotetraploids, respectively, formed between maize and *Z. perennis* by natural genome extraction. **(a-c)** show means with a standard deviation bar; **(a1 – c1)** shows the median with upper and lower quartiles. Different small letters (a, b, and c) represent significant statistical differences among means (one-way ANOVA, followed by Tukey test) between different groups for respective traits, and colored filled circles show individual values of a member within the respective population. **(d-f)** represent means with standard deviations of seeds, number, seeds setting (%), and pollen viability of different populations of Amphitetraploid hybrids. (**g-i**) shows days to initiation of tasseling, silking, and anthesis of the first three selfed generations of two independent populations of amphitetraploid maize. Different small letters (a, b, and c) represent significant statistical differences among means (one-way ANOVA, followed by Tukey test) between different groups for individual traits, and colored filled circles show individual values of members within a respective population.

### Transgenerational chromosome dynamics in nascent near-allotetraploids

In a cohort of 300 plants of S2-S6, mostly observed individuals were 2n=40 (35.66%, 107 individuals), followed by 2n=39, 41, 42, and 38 found in 77 (25.66%), 55 (19.00 %), 28 (10.00%) and 19 (6.33%) plants, respectively, while <38 (minimum 37) and >42 (maximum 45) were very rare (**Table 1**). Although chromosome distribution within the lines across the generations was variable (range, 2*n* = 37-45), but mean chromosome number was preferably maintained at approximately tetraploid level 2n=40 (**Table S3**). There was no significant difference among S2, S4, S5, and S6 in terms of chromosome number (ANOVA, *P* > 0.05), while S3 had a significantly lower mean chromosome number of 39.54 from all other generations (ANOVA, *P* = 0.0057). The mean chromosome number in S3 (39.54) was also significantly different from 40 (one-sample *t*-test, *P* < 0.05.); however, the distribution of chromosomes was not normal and was skewed toward the accumulation of more chromosomes (Shapiro-Wilk test, W= 0.890; *P* < 0.05; Table S3; skewness of 0.466). In all remaining generations (S2, S4, S5, S6), the mean chromosome number was not significantly different from 40 (one-sample *t*-test, *P* > 0.05); however, distribution was not normal and skewed towards the accumulation of more chromosomes (**Table S3**). Somatic chromosome instability was not observed within the cells of a given individual, indicating normal mitosis there.

**Table 1.**
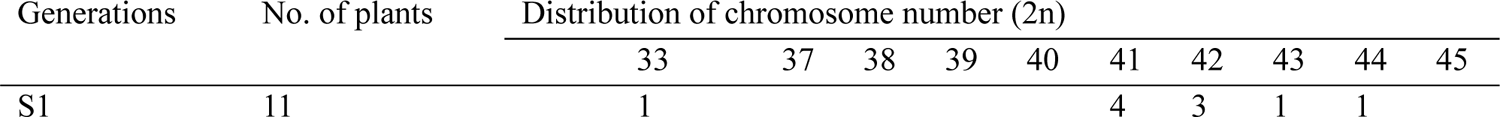
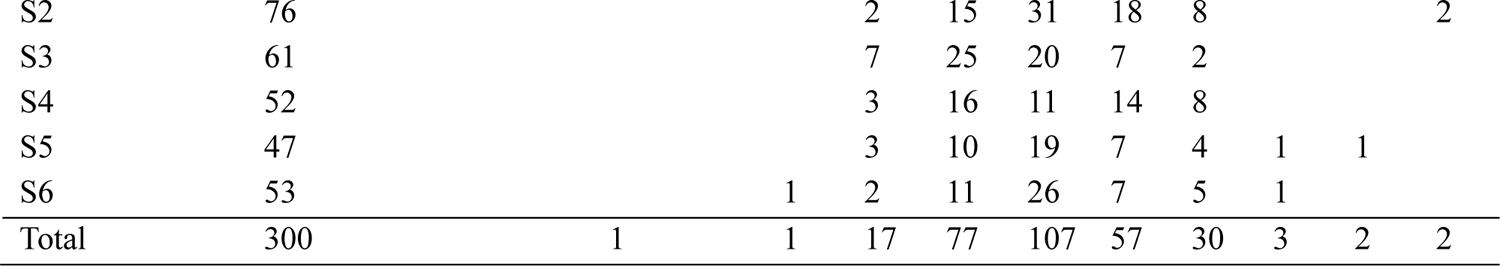
Summary of somatic chromosomes distribution in the populations of nascent near-allotetraploids (S1–S6)

### The stability of subgenome chromosomes was different for three parental genomes in selfed generations (S1-S6)

Multi-color GISH analysis could differentiate the chromosomes complement of three genomes and detect large-scale chromosome changes (recombinant chromosomes/translocation) among subgenomes in a cell. Mc-GISH results per individual of S1-S6 generations are presented in **Table 2 and Figure S6-S7)**. Ten analyzed plants of the first selfed generation (S1) harbored maize (M), *Z. perennis* (P), and *T.dactyloides* (T) chromosomes in the range of 23-28, 5-17, and 1-5 (Mean 25.20M+13.3P+2.50T), respectively (**Figure S6a-c**). A drastic loss occurred for *Tripsacum* chromosomes, while the M and P genomes were stably inherited relative to the immediate parent (**Table 2**). The 36 analyzed plants of the second selfed generation S2 had M, P, and T chromosomes in the range of 21-32, 10-19, and 0-2 (Mean 25.04M+14.51P+0.53T), respectively (**Figure S6d-f**); of them, 20 plants (55.6%) had eliminated all *Tripsacum* chromosomes, 13 plants (36.1%) harbored 1 and 3 plants (8.3%) contained 2 *Tripsacum* chromosomes (**Table S2**). In 29 observed plants of the S3 generation, M, P, and T chromosomes ranged from 22-28, 11-17, and 0-2 (Mean 25.03M+14.07P+0.21T), respectively (**Figure S6g-i**). Of them, 82.8% had eliminated all *Tripsacum* chromosomes, 13.8% had 1, and 3.4% harbored 2 *Tripsacum* chromosomes (**Table S2**). The 16 investigated individuals of the S4 generation harbored M, P, and T chromosomes in the range of 24-30, 12-16, and 0-1 (Mean 26.06M+13.94P+0.06T), respectively (**Figure S6k-m**). Of them, 15 (93.7%) had eliminated all *Tripsacum* chromosomes, while only 1 (6.3%) had 1 *Tripsacum* chromosome (**Table S2**). In 33 analyzed plants of the S5 generation, M, P, and T chromosomes ranged from 22-30, 10-18, and 0-1 (Mean 25.94M+14.0P+0.03T), respectively (**Figure S6n-p)**. Of them, only one plant had 1 *Tripsacum* chromosome, and all remaining individual had eliminated all *Tripsacum* chromosomes (**Table S2**). In 16 analyzed individuals of the S6 generation, M and P chromosomes ranged from 24-28 and 12-16 (Mean 26.06M+14.00P), respectively; and no individual was detected carrying *Tripsacum* chromosomes (**Table S2**). The selfed generations (S2-S6) were not significantly different in terms of maize and *Z. perennis* chromosomes distribution (ANOVA, *P* > 0.05), and variance decreased by advancing the generations (**Table 2**). The accumulative results of S1 to S6 generation indicated that *T. dactyloides* chromosomes were eliminated, while maize and *Z. perennis* belong to the same genus and so their chromosomes may pair with each other being relatively stably transmitted to the next generations with minor numerical changes possibly for genome balance. Based on M. P and T chromosomes distribution within a given genome, there were 10, 26, 16, 10, 21, and 10 kinds of cytotypes in S1, S2, S3, S4, S5, and S6 generations populations, respectively, with the major observed cytotypes of 24-26M+14-16P chromosomes (**Figure S7**), indicating a higher genomes heterogeneity at the chromosome level. There was no significant correlation between chromosome number or chromosome rearrangements detected by GISH (*P* > 0.05) with pollen viability or seed fertility. Nevertheless, a significant negative correlation existed between pollen viability and maize chromosomes (r = −0.7787, *P* = 0.0391), and vice versa, there was a positive correlation between pollen viability and *Z. perennis* chromosomes (r = 0.8036, *P* = 0.0294).

**Table 2.** Summary of subgenomes chromosome inheritance from S1 to S6 generation of nascent-near allotetraploids

| Generation | N | Maize |  |  | *Z. perennis* |  |  | *T. dactyloides* |  |  |
| --- | --- | --- | --- | --- | --- | --- | --- | --- | --- | --- |
| | | Mean ± SD | Variance | Range | Mean ± SD | Variance | Range | Mean ± SD | Variance | Range |
| S1 | 10 | 25.20 ± 1.55 | 2.40 | 23-28 | 13.3 ± 3.33 | 11.12 | 5-17 | 2.50 ± 1.27 | 1.611 | 1-5 |
| S2 | 36 | 25.04 ± 2.52 | 6.36 | 21-32 | 14.51 ± 1.88 | 3.55 | 10-18 | 0.53 ± 0.65 | 0.43 | 0-2 |
| S3 | 29 | 25.03 ± 1.57 | 2.46 | 22-28 | 14.07 ± 1.81 | 3.28 | 11.17 | 0.21 ± 0.49 | 0.24 | 0-1 |
| S4 | 16 | 26.06 ± 1.98 | 3.93 | 24-30 | 13.94 ± 1.24 | 1.53 | 12-16 | 0.06 ± 0.25 | 0.06 | 0-1 |
| S5 | 33 | 25.94 ± 1.97 | 3.87 | 22-30 | 14.00 ± 1.95 | 3.81 | 10-18 | 0.03 ± 0.17 | 0.03 | 0-1 |
| S6 | 16 | 26.06 ± 1.24 | 1.53 | 24-28 | 14.00 ± 1.55 | 1.33 | 12-16 | 0.00 | - | - |
| Total | 140 | 25.53 ± 2.13 | - | 21-32 | 14.14 ± 2.06 | - | 5-23 | 0.37 ± .082 | - | 0-5 |

### Karyotype stability was different among individual chromosomes of subgenomes

By observing extensive numerical variations in the chromosomes of M and P genomes with a trend of a total chromosome number (2n) around 40. A pertinent question arose: do all chromosomes of M and P genomes show similar propensities for whole chromosome loss or gain? Individuals with clear chromosome spreads were karyotyped (**Figure 4, Table 3 and Table S4**). Subgenomic chromosomes were markedly different for gain and loss. Among ten maize chromosomes, M5, M6, and M7 showed higher frequencies of chromosome gain than the rest of the chromosomes. All chromosomes except M3 were involved in loss/gain. Three maize chromosomes, M1, M2, M4, and M10, were monosomic in 12 (35%), 2 (5.88%), 2 (5.88%), and 6 (17.64%) individuals, respectively. Chromosome M7 generated extra copies in 58.33% of plants, while M5 and M6 were hyperploid in 83.33% of individuals. Chromosomes M3, M5, M6, M7, M8, and M9 were not lost in any analyzed plant, probably due to the detrimental effect of the combination of monosomy of either of these five maize chromosomes and the chromosome gain/loss of other chromosomes in maize or homologous chromosomes of *Z. perennis*. According to Gene Balance Hypothesis, dosage-sensitive genes are more likely associated with protein complexes, e.g., regulatory complexes. In a simple monosomy case (only lost one certain chromosome), changing the protein level of one subunit (a dosage-sensitive gene on this chromosome) to 1/2 (monosomy), the stoichiometry of the whole functional complex is changed and thus has detrimental effects but plant still show survival. But for the aneuploid combination example, if the dose of other subunits (subunits) on the different chromosomes is disrupted (monosomy, trisomy, or tetrasomy) in the same cell, you would expect the complex is affected to a greater extent, which might be lethal and thus selected against. For *Z. perennis,* monosomy and nullisomy were prevalent states; chromosomes P3, P13, and P16 were often nullisomic, while P7 and P15 generated extra copies more often. A lost copy of *Z. perennis* was probably compensated by a copy from the maize genome to maintain genome balance. A total of 346 chromosomes were lost compared to 322 gained chromosomes in all 48 analyzed plants.

**Figure 4.**
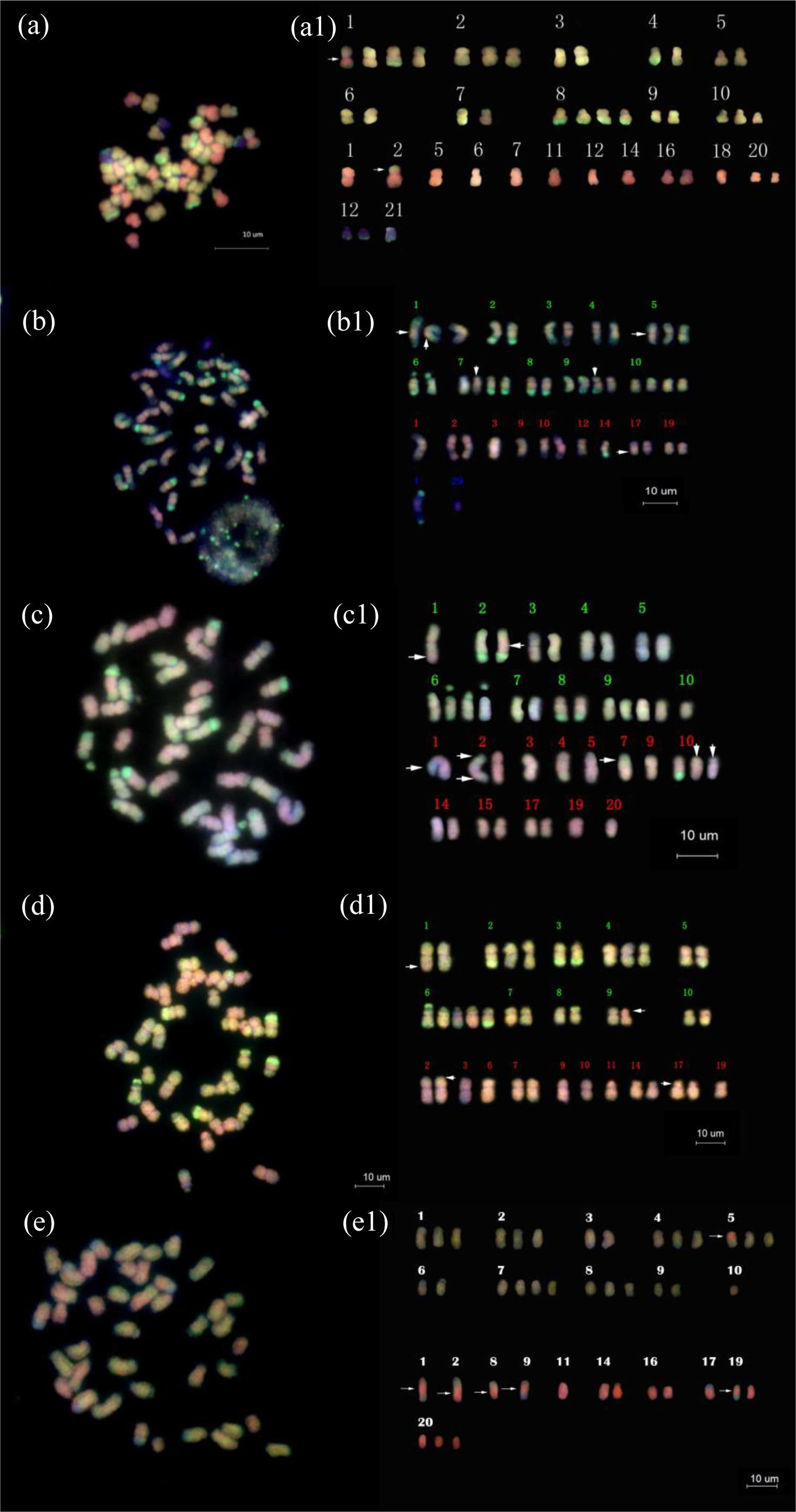
Mitotic karyotype of nascent near-allotetraploids (NsAt) at different generations and homoploid recombinational allotetraploid hybrids (RcAt). GISH was carried out with total genomic DNA probes of *Z. mays* ssp. *mays* (green) and *Z. perennis* (red), while *T. dactyloides* chromosomes were recognized by DAPI (blue) from merged pictures. Arrows indicate the position of translocation/breakpoint. From (**a1)** to (**e1),** the upper two rows display maize chromosomes, while the lower 1-2 rows show *Z. perennis* chromosomes. (**a-a1)**, (**b-b1), (c-c1)** and (**d-d1)** and **(e-e1)** display genome complements and karyotype of S1-7 (2n=42), S1-4 (2n=43), S2-14-13 (2n=41), S6-4-5-1-4-5-2 (39), and S6-14-2-9-2-6-12-2n= 41), respectively.

**Table 3.** Distribution of the number of plants showing variables (from 0 to 4) of each chromosome of maize (M) and *Z. perennis* (P)

|  | Nullisomy | Monosomy | Trisomy | Tetrasomy | Total |
| --- | --- | --- | --- | --- | --- |
| M1 | 0 | 12 | 4 | 10 | 26 |
| M2 | 0 | 2 | 6 | 6 | 14 |
| M3 | 0 | 0 | 0 | 0 | 0 |
| M4 | 0 | 2 | 8 | 0 | 10 |
| M5 | 0 | 0 | 18 | 22 | 40 |
| M6 | 0 | 0 | 14 | 26 | 40 |
| M7 | 0 | 0 | 6 | 22 | 28 |
| M8 | 0 | 0 | 2 | 10 | 12 |
| M9 | 0 | 0 | 6 | 8 | 14 |
| M10 | 0 | 6 | 4 | 2 | 12 |
| Total | 0 | 22 | 68 | 106 | 392 |
| P1 | 30 | 14 | 0 | 0 | 44 |
| P2 | 26 | 12 | 4 | 0 | 42 |
| P3 | 36 | 8 | 0 | 0 | 44 |
| P4 | 20 | 26 | 0 | 0 | 46 |
| P5 | 28 | 18 | 0 | 0 | 46 |
| P6 | 18 | 16 | 0 | 0 | 34 |
| P7 | 9 | 11 | 8 | 16 | 44 |
| P8 | 30 | 16 | 0 | 0 | 46 |
| P9 | 10 | 20 | 4 | 4 | 38 |
| P10 | 16 | 16 | 2 | 0 | 34 |
| P11 | 26 | 20 | 0 | 0 | 46 |
| P12 | 34 | 8 | 2 | 0 | 44 |
| P13 | 38 | 10 | 0 | 0 | 48 |
| P14 | 18 | 15 | 0 | 4 | 37 |
| P15 | 24 | 12 | 0 | 0 | 36 |
| P16 | 36 | 8 | 0 | 0 | 44 |
| P17 | 24 | 18 | 0 | 0 | 42 |
| P18 | 26 | 12 | 0 | 0 | 38 |
| P19 | 12 | 26 | 0 | 0 | 38 |
| P20 | 26 | 12 | 2 | 0 | 40 |
| Total | 487 | 298 | 22 | 24 | 1662 |

### Structural chromosome variations (SCV) detected by mc-GISH were predominantly between homoeologous chromosomes

Large-scale structural chromosome variations (SCV) were detected in the first six generations of nascent near-allotetraploids by mc-GISH with the results are presented in **Table 4 and Table S5**. Of 139 analyzed plants, 137 (98.56%) had chromosome translocations between maize and *Z. perennis* chromosomes (MP), ranging from 1-10, with an average of 4.05/plant, but there was no significant difference among selfed generations in terms of translocated chromosomes (ANOVA, *P* = 0.685). In 137 plants (S1-S6), mostly observed chromosomes translocations/plant were 3 observed in 43 individuals (31.39%), followed by 4, 6, 5, and 2 translocations detected in 35 (25.55%), 17 (12.41 %), 14 (10.22) and 14 (10.22%) plants, respectively, while <2 (minimum 1) and >6 (maximum 10) were very rare, indicating that highly rearranged genomes might be selected against, and did not transmit to the next generation. The differences in translocated chromosomes were observed between the plant of the same generation and between the generations. Still, various plant cells of an individual carried similar translocations, suggesting only meiosis produced chromosomal rearrangements. Careful visual observation of intergenomic translocations reflected that all translocations were reciprocal or non-reciprocal exchanges, most probably among homoeologous chromosomes. In most observed plants (40.74%), maize chromosome M1 showed reciprocal homoeologous translocation with P1 or P2. Further, homoeologous pairs M5/P10 and M7/P15 were involved in homoeologous recombination in 22.22% and 13.5% of analyzed plants, while all maize chromosomes except M6 showed non-reciprocal chromosome exchanges at least once (**Table S5**). Transgenerational karyotype analysis was carried out using individuals descended from a single plant for observing inheritance and stability of translocated chromosomes across selfed generations. The translocated chromosomes detected in selected individual of the S3 was not found in the progeny of the same individual. Additionally, chromosome copy number deviation was higher in chromosome pairs harboring translocated chromosomes, indicating that highly rearranged chromosomes might not transmit to the next generation. A normal chromosome generated an additional chromosome copy for balancing gene dosage. The chromosome translocations observed in immediate parents did not match precisely with progeny, suggesting that the generation of chromosomal rearrangements was an ongoing process, but most of the rearranged chromosomes selected against and only favoring translocation may persist and retain.

**Table 4.**
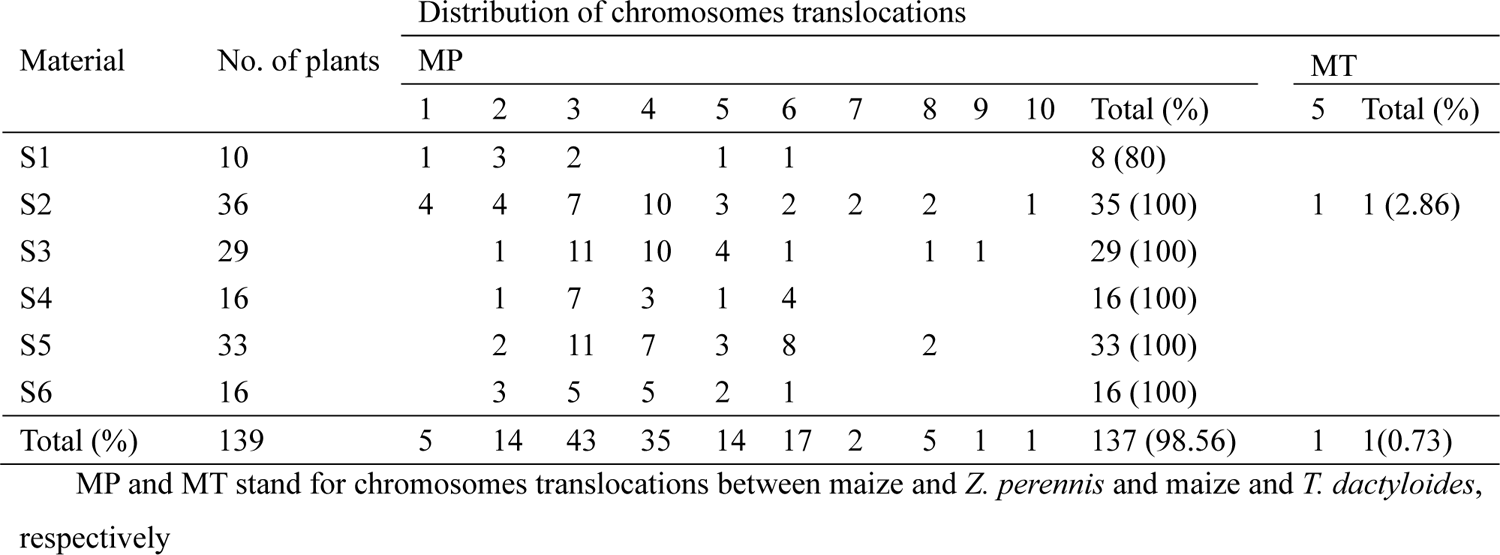
Large-scale SCV distribution in analyzed populations of the generations (S1-S6) of nascent-near allotetraploids

### Structural chromosomes variations detected by 45S and 5S rDNA loci probes

Maize, *Z. perennis,* and *T. dactyloides* had two, four, and six 45S and 5S gene loci, respectively (**Figure S8**). The signal patterns of 45S and 5S loci are different in size and location, thus allowing effective identification. The MTP had six 45S loci and six 5S loci (**Figure S9a and Figure S12a**), with no apparent loss relative to parents. In the S2 population, three (30%) plants changed 45S, and 1 (10%) changed 5S loci number relative to parental additive number of 4 (**Figure S10 and Table S6**). In the S6 population, 1 (3.8%) individual changed 45S, and 2 (16.66%) changed 5S loci number relative to the parents (**Figure S9 and Table S6**). Despite loci number changes, other commonly observed changes were small chromosome fragments with or without 45S loci but without CentC signals (**Figure S10a, e and f)**, which were broken chromosomal fragments produced as a result of chromosome rearrangements or mechanical force during FISH slides preparations. Plants with two or three 45S loci show loss of respective chromosomes or segments. Chromosomes of maize and *Z. perennis* bearing 45S loci also harbored nuclear organizer region (NOR) and secondary constriction. In this study, the chromosome detected with secondary constriction but without 45S signals (**Figure S10a**) or chromosomes bearing 45S loci but without the obvious secondary construction were detected (**Figure S10c**), indicating that the respective chromosome might have lost 45S loci or copy number decreased to a level that did not fulfill identification requirements. A line (S6-14) showed a chromosome pair bearing one 45S loci with secondary constriction, while the second pair had 45S loci, but did not show secondary constriction (**Figure S9f**). One family of S6 generation (S6-31-14-2-9-2-6) has changed the 5S loci position from the distal end of a long arm to a position to a centromere with a reduced 5S loci size (**Figure S13K and L**), which indicates the breakage and fusion of 5S loci. A plant of the S6 generation was detected bearing eight 45S loci with 84 chromosomes (**Figure S11a1-a3**), probably due to genome doubling in root, but the plant died before reaching maturity, so it could not be investigated further.

### AFLP analysis revealed stable introgression from *T. dactyloides*

GISH could effectively differentiate sub-genome chromosomes and detect large-scale chromosome translocation but cannot reveal small-scale genetic exchanges. Therefore, we used the AFLP markers due to their genome-wide coverage, no prior need for genomic/genetic knowledge, and their capacity to generate many DNA fragments. The results are presented in **Figure S14**. All analyzed individuals of the 6^th^ selfed generation had eliminated all *Tripsacum* chromosomes, so it was reasonable to detect small-scale genetic transfer from *Tripsacum,* if any kind, by using AFLP. The AFLP fragments were divided into seven classes: shared DNA fragments between three parents (Mz+Zp+Tr), shared between either of two parents (Mz+ZP: Mz+Tr: Zp+Tr), and unique parental DNA fragments (Mz: Zp: Tr). All three analyzed lines of NsAt, even at the S6 generation, showed the presence of 6 to 8% unique *Tripsacum* DNA (Tr) fragments, indicating stable introgression and transmission of *T. dactyloides* genetic material in these allotetraploid populations. In all studied materials, the variation level of the three parental bands differed, and the most drastic changes occurred just after hybridization or before the second selfed generation (S2). However, genomic changes took place comparatively slower in the following generations. The proportion of *Tripsacum-*specific fragment loss was maximum (> 80%), followed by shared sequences between *Z. perennis* and *T. dactyloides* (Mz+Tr), while minimizing loss of genome-specific sequences was found for maize. The shared sequences between three parents (Mz+Zp+Tr) were comparatively more stably transferred in succeeding selfed generations as their loss was minimum (∼10%) that was followed by shared sequences between maize and *Z. perennis* (∼ 16%). Three studied sister lines showed a similar pattern of fragment loss or retention, indicating observed genomic variations were reproducible and non-random to some extent. Moreover, the relative abundance of maize and *Z. perennis* fragments than *T. dactyloides* reflected their closer genetic relationship as both belong to the same genus *Zea*; while *Tripsacum* diverged from *Zea* a few million years ago (Welker et al. 2020), although they all share a polyploidy event (Estep et al. 2014). Therefore, similarities in AFLPs have the potential to be coincidental, given their nature.

### Construction of autotetraploid maize genomes for improved fertility and bivalent pairing

Developing true-breeding allotetraploid maize has long been one of the important goals of polyploidy maize breeding. In an attempt to utilize nascent near-allotetraploids as a genetic bridge to introduce differentiated chromosomes and to induce chromosomes rearrangement for restructuring the genome of tetraploid maize, so that differentiated chromosomes do not pair with normal maize chromosomes, we hybridized selected individuals of the S6 population with 2 autotetraploid maize inbreds P1-vv (abbreviated as BP) and TWF9 (abbreviated as BT), and followed by three selfed generations. A remarkable improvement in seed set and pollen viability relative to immediate parents was observed in these populations, which was maintained for at least up to the F3 (**Figure 2q and Figure 3d-f**). Plant exhibited synchronized tasseling, silking, and anthesis well and showed fast growth earlier than cultivated maize (**Figure 3g-i**). It was important to examine the genomes of constructed allotetraploid maize for karyotype uniformity and stability. Therefore, we analyzed transgenerational chromosome inheritance patterns (**Table 5 and Table S7**). Notably, no chromosome instability within an individual’s somatic cells was detected, reflecting normal mitosis. Any numerical or structural chromosome variation is rooted in abnormal meiosis. In all populations (F1-F3), the mean chromosome number did not significantly change from 40 (one-sample *t*-test, *P* > 0.05). The frequency of aneuploidy (missing or extra chromosomes from the theoretical tetraploid value (2n=40) significantly decreased from 60% for BPF1 and 66.7% for BTF1 to 21.3% in BPF3 and 20% in BTF3, respectively. There was almost the same trend for BPT2, with slightly higher aneuploidy in BTF3 (21.3%) than in BTF2 (14.3%), while the mean chromosome number remained approximately 40 and variance decreased by advancing the generations (**Table 5**). In a cohort of 171 plants of F1-F3, mostly observed individuals were 2n=40 (69.01%, 118 individuals), followed by 2n=41, 42, and 39 found in 16 (9.36%), 16 (9.36 %), and 15 (8.77%) plants, respectively, while <39 (minimum 38) and >42 (maximum 43) were very rare (**Table S7**).

**Table 5.** Statistics of total chromosome number (2n) in the selfed generations (S1-S6) of nascent-near allotetraploids

| Population | N | Mean | Variance | Range | Aneuploidy % |
| --- | --- | --- | --- | --- | --- |
| BPF1 | 20 | 40.05 ± 1.19 | 1.4184 | 38-42 | 60 |
| BTF1 | 12 | 40.25 ± 1.21 | 1.4773 | 38-42 | 66.7 |
| BPF2 | 37 | 40.324 ± 0.82 | 0.6697 | 39-42 | 35.1 |
| BTF2 | 21 | 40.095 ± 0.54 | 0.2905 | 39-42 | 14.3 |
| BPF3 | 47 | 40.191 ± 0.71 | 0.506 | 39-42 | 21.3 |

### GISH analysis of tetraploid maize hybrids and progeny revealed genome retention bias and extensive genome restructuring

From the F1 to F3, most of the *Z. perennis* chromosomes are lost or present in the form of translocated chromosomes. Simultaneously, the loss of *Z. perennis* chromosomes is compensated by extra copies of maize chromosomes to maintain the genome at/near the tetraploid level. The detailed per plant results of mc-GISH are presented in **Table 6** and **Figure S15**. Based on the proportion of maize (M) and *Z. perennis* (P) chromosomes and translocated chromosomes between maize and *Z. perennis* genomes (MP), there were 16 different cytotypes in 16 BPF1 plants identified by mc-GISH (**Figure S15a**). In the BPF1 population, M, P, and MP chromosomes ranged from 25–37, 3-10, and 0-7 (average, 32.21, 5.78, and 2.43), respectively (**Figure S16a-c**). In 12 BTF1 plants, M, P, and MP chromosomes ranged from 26–36, 3-8, and 0-5 (average, 32.58, 5.41, and 2.25), respectively (**Figure S16d-f)**, with 10 different cytotypes (**Figure S15b)**. There was no significant difference in chromosome complements between BPF1 and BTF1 (ANOVA, *P > 0.05*). Thirty-six BPF2 plants harbored M, P, and MP chromosomes in the range of 27-39, 0-9, and 0-7 (average, 34.27, 2.88, and 3.38), respectively (**Figure S16g-i)**. The M, P, and MP chromosomes in twenty BTF2 ranged from 30-37, 1-4, and 2-7, respectively (**Figure S16k-m)**. BPF2 and BTF2 populations were not significantly different regarding chromosome complements (ANOVA, *P > 0.05*). Forty-two BPF3 plants identified by GISH showed 21 different cytotypes (**Figure S15e)**, in which M, P, and MP chromosomes ranged from 32-39, 0-3, and 2-7 (average, 35.39, 1.02, and 3.83), respectively (**Figure 5a-f**). Thirty-six BTF3 plants formed 18 different chromosome complements (**Figure S15f)**, in which M, P, and MP chromosomes ranged from 31-40, 0-2, and 2-10 (average, 35.47, 0.86, and 3.75), respectively (**Figure 5g-m**). There was no significant difference in both F3 populations regarding chromosome complements (ANOVA, *P > 0.05*). Considering BP and BT populations as one group, the mean for M chromosomes significantly increased from 31.47 - 34.25 - 35.43 in the F1 - F2 - F3, respectively (ANOVA, *P < 0.0001*). Meanwhile, the means for the P chromosomes significantly decreased from 6.76 - 3.14 - 0.95 in F1 - F2 - F3, respectively (ANOVA, *P < 0.0001*). The mean for structural chromosome variation detected by GISH significantly increased from 2.43 - 3.48 - 3.80 in F1 - F2 - F3 generations, respectively (ANOVA, *P = 0.0005*).

**Figure 5.**
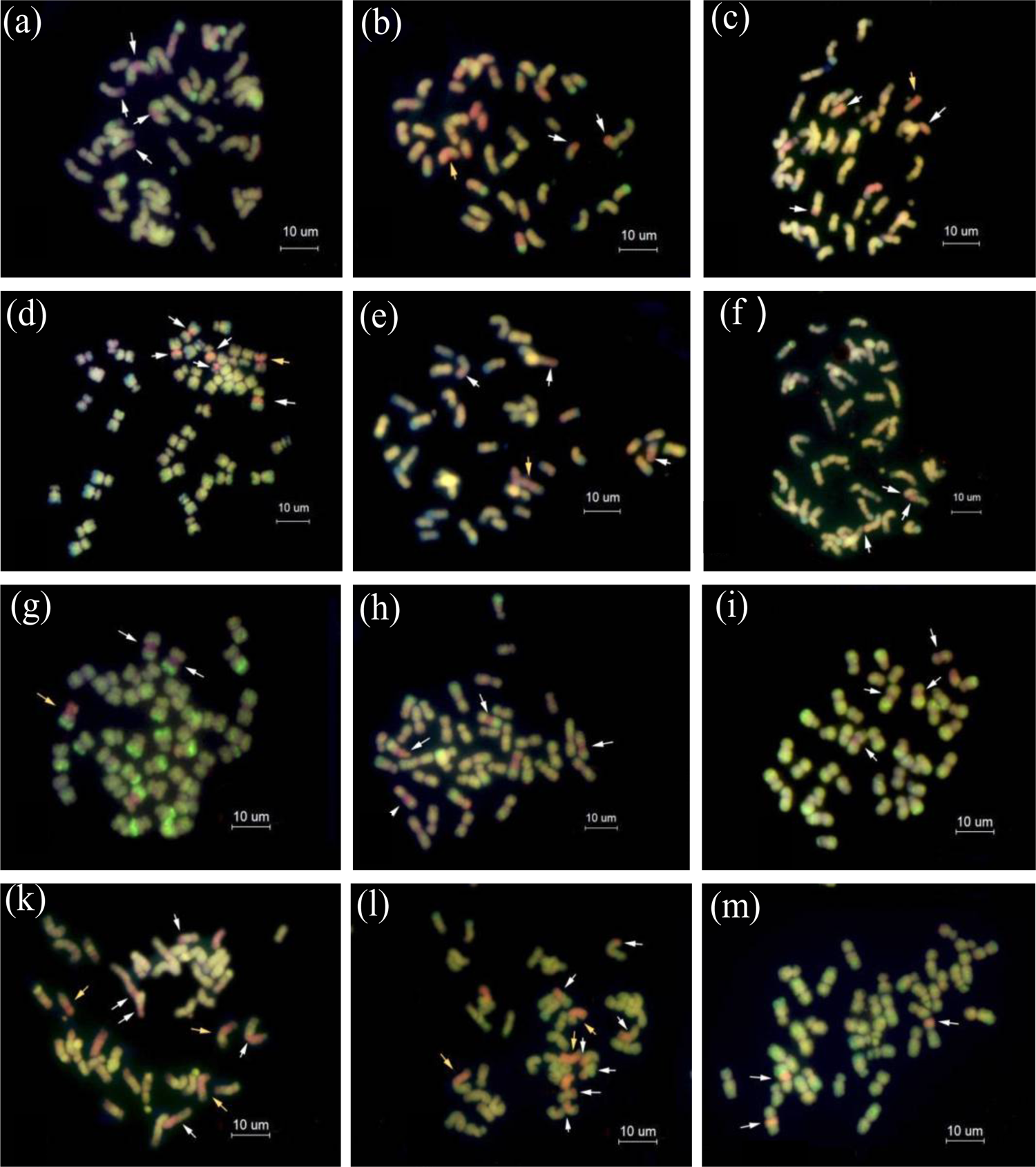
mc-GISH analysis of root tip mitotic chromosomes of BP and BT populations of the F3 generation. Here, (**a)**:18-49-3-1; (**b)**: 18-52-10-5; (**c)**:18-52-1-1; (**d)**: 18-52-2-5; (**e)**:18-52-10-4; (**f)**: 18-50-7-4 plants of BP-F3 population. (**g)**: 004-4-3; (**h)**: 006-1-1; **(i)**: 006-1-2; (**k)**: 006-3-2; (**l)**: 006-3-3; (**m)**: 014-1-4. Green and pink colors show the chromosomes of maize (M) and *Z. perennis,* respectively. White arrows show translocations between maize and *Z. perennis* chromosomes (MP), and yellow arrows point out translocations between *Z. perennis* and maize (PM). The bar is 10 um

**Table 6.**
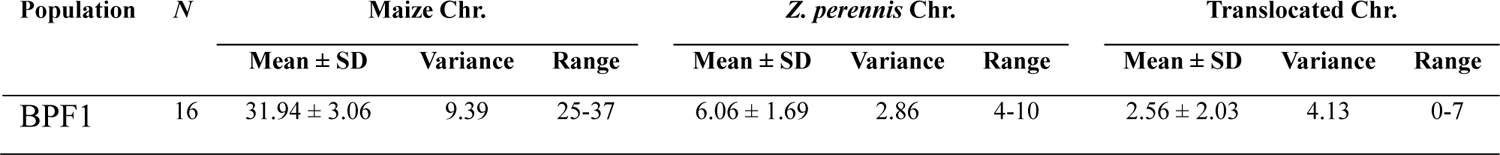
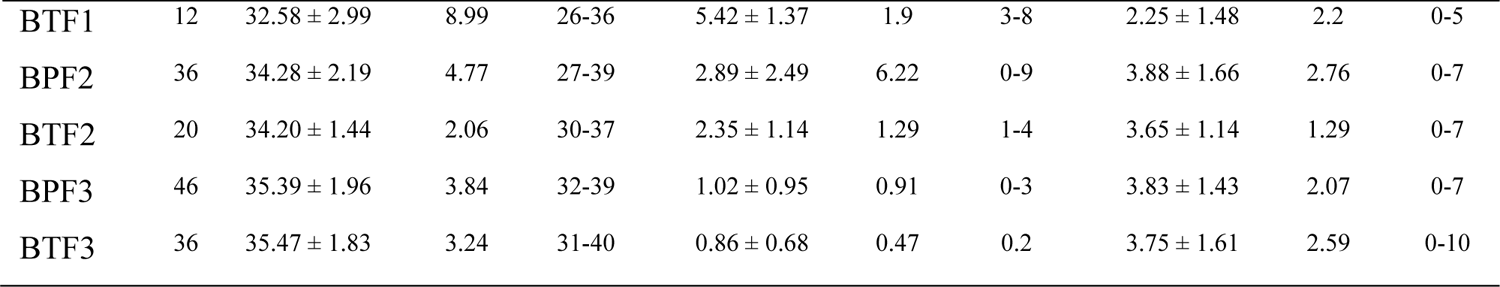
Summary of sub-genomes chromosome inheritance in selfed generations of reconstructed allotetraploid maize from F1 to F3 generations

Karyotype analysis of 14 individuals from F2 and their progenies showed the presence of *Z. perennis* chromosomes 3, 6, 8, 12, 13, 17, and 19, of which chromosome transmission rates of chromosomes 3, 8, 12, and 13 were higher. Meanwhile, maize chromosomes 1, 2, 3, 4, 7, 8, 9, and 10 showed chromosome translocations with the P genome, of which M2, M4, and M8 showed stable transmission of translocations to the next generation, indicating stable integration of P genome genetic material into the M genome (**Figure 6**).

**Figure 6.**
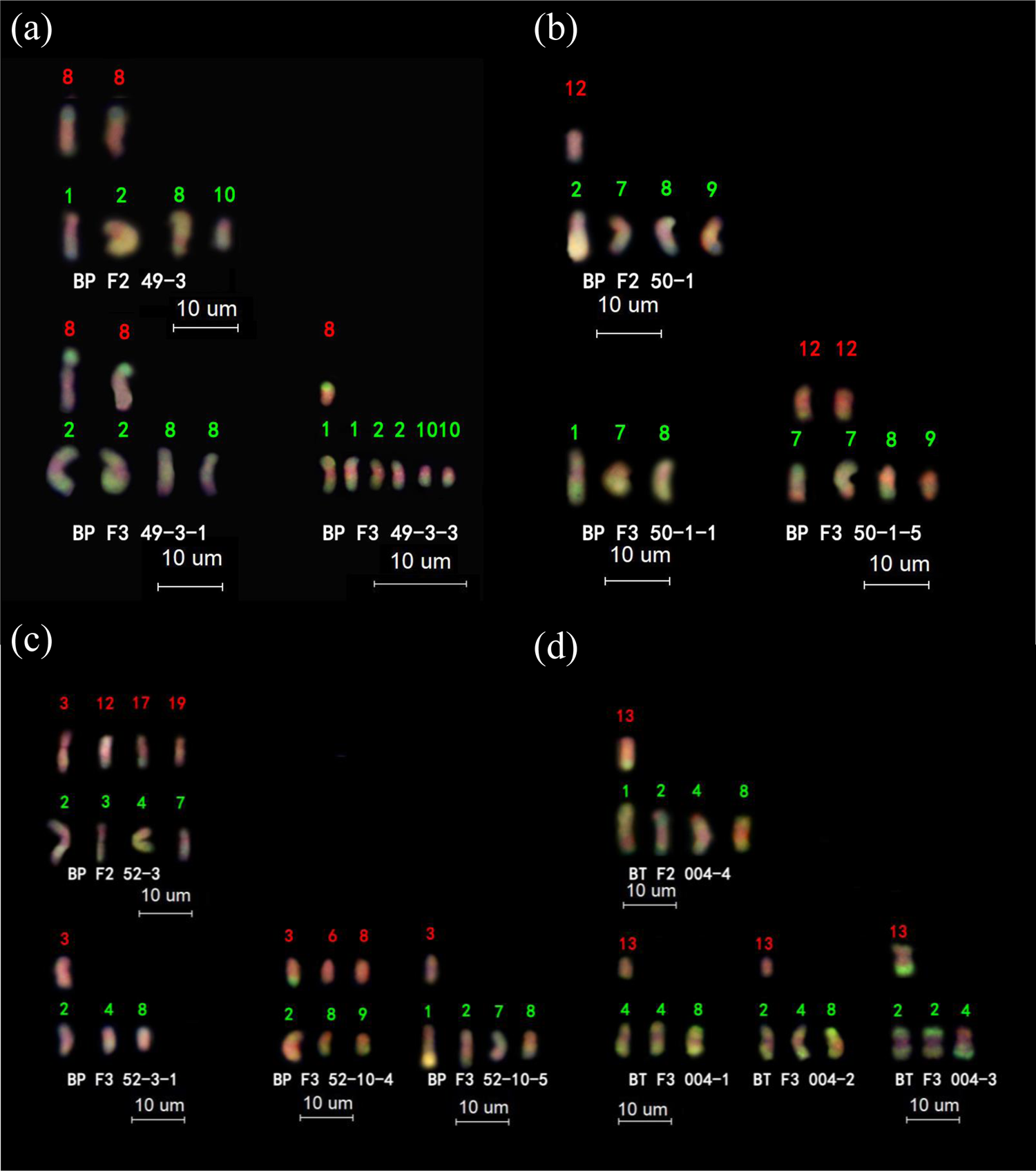
Karyotype analysis of exogenous genetic material transfer and transmission in F2:F3 generations of amphitetraploid maize hybrids. GISH was carried out with total genomic DNA probes of *Z. mays* ssp. *mays* (green) and *Z. perennis* (red), while chromosomes were stained by DAPI (blue). Only chromosomes from *Z. perennis* genome and translocated chromosomes are shown. Green or red digits over chromosomes represent the respective chromosome numbers of maize and *Z. perennis*, respectively. White alphabetic and digits show generation and plant codes, respectively. In (**a-d)**, the upper layer displays foreign chromosomes/fragments in a tetraploid maize genome, while the lower layer shows their inheritance in immediate progenies. The bar is 10 um

### Meiotic behavior of constructed tetraploid maize showed chromosomes of maize and *Z. perennis* have merged

Meiotic behavior of 43 individuals (23 from BPF3 and 20 from BTF3) was observed along with parents by analyzing at least 15 cells/individual (**Table S8**). The frequency of abnormal meiosis (presence of laggard chromosomes or chromosome bridges) was 52.38% in S6 generation, 63.26% in P1-vv, and 49.73% in Twf9, which was significantly higher from BPF3 (23.26%) and BTF3 (26.9%) population (ANOVA, *P > 0.005).* The average meiotic configuration in BPF3 (five plant analyzed, 2n=40) was 0.11 univalent (Ⅰ) + 8.95 bivalent (Ⅱ) + 0.41 trivalent (Ⅲ) + 5.19 tetravalent (Ⅳ), with a range of 0-0.27I + 8.4 – 9.33II + 0.20-0.67III + 5.00 – 5.60IV (**Table S9**). The average meiotic configuration in BTF3 (five plants analyzed, 2n=40) was 0.19Ⅰ + 8.33Ⅱ + 0.57Ⅲ + 5.36Ⅳ, with a range of 0.13 – 0.20I + 8.07 – 8.60II + 0.33 – 0.97III + 5.07 – 5.67IV (**Table S10**). Both populations showed significant improvement over bivalent pairing and reduction in multivalent formations relative to autotetraploid maize parent P1-vv (Average; 0.32Ⅰ + 3.40Ⅱ + 0.20Ⅲ + 8.07Ⅳ), and Twf9 (Average; 0.26Ⅰ + 3.27Ⅱ + 0.40Ⅲ + 8.00Ⅳ). From the F3, 77 meiotic cells were analyzed by mc-GISH; the main findings are: (1) chromosomes of *Z. perennis* without structural variations paired preferentially; (2) Chromosomes with structural variation paired with each other or with normal maize chromosomes to produce bivalents or a tetravalent; (3) univalents from the *Z. perennis* genome were not found (**Figure 7**). The results indicated that genomes of the constructed maize hybrids had merged maize and *Z. perennis* chromosomes, which suggested that re-domestication of tetraploid maize could be accomplished by allopolyploidization and subsequent sub-genomes extraction. It can develop diploid-like meiotic behavior with gradual improvement.

**Figure 7.**
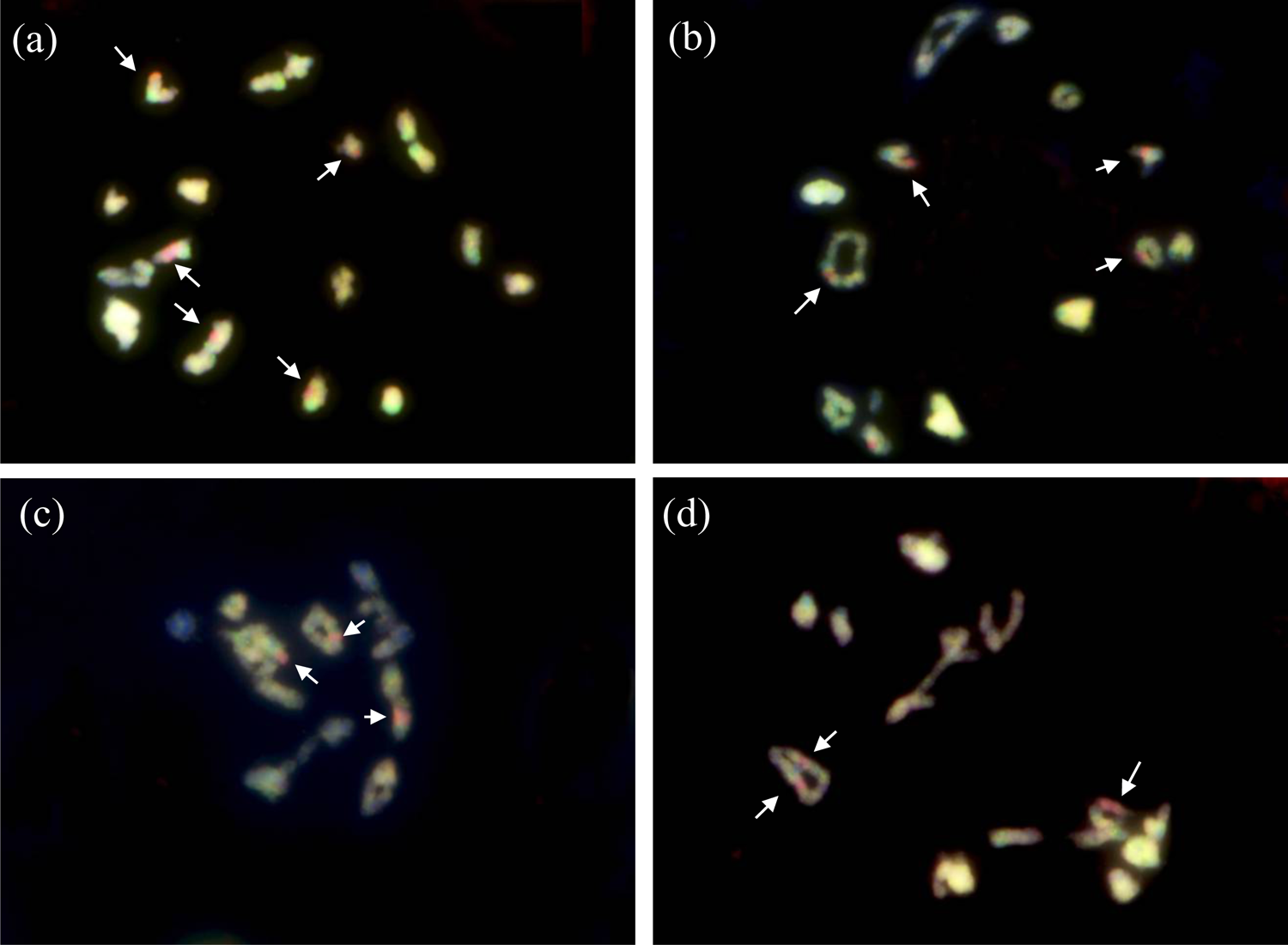
Meiotic chromosome behavior at diakinesis in PMC of amphitetraploid maize hybrids. (**a)** to (**d)** are examples showing chromosome pairing, that is, chromosomes from *Z. perennis* genome without structural variations preferentially paired; (2) chromosomes with structural variation paired with each other or with normal maize chromosomes to produce bivalent or tetravalent; (3) univalent from *Z. perennis* genome were not found. The arrows direct chromosomes or translocated chromosome segments from *Z. perennis*. (**a)** to (**d)** display genomes of BPF3-49-3-1, BPF3-51-4-3, BPF3-50-1-1, and BPF3-51-4-1, respectively.

## Discussion

### Synthetic allopolyploids are model materials for studying hybrid speciation among polyploid species

The creation of polyploids and their genetic breeding has long been one of the important means to increase genetic variations and trait improvement (Mason and Batley 2015; Udall and Wendel 2006). If polyploid progenies of desired traits are successfully created by distant hybridization or using physical and chemical means, and they are sexually fertile, new lines with steadily stable genetic variation can be formed through self-breeding, which plays an important role in the study of the polyploids synthesis, biological evolution, heredity, and breeding. Existing polyploid species have developed through the long-run process of genome diploidization and functional coordination (Soltis et al. 2009; Van de Peer et al. 2009; Wendel 2000). However, when different species or different genomes are combined, duplicated, extracted, and substituted to create new polyploid materials, they show poor early adaptation (Chester et al. 2012; Matsushita et al. 2012; Pontes et al. 2004; Zhang et al. 2013).

The process of plant genome evolution after polyploidy is dynamic. Polyploidy generally produces phenotypic and karyotypic heterogeneity (Gou et al. 2018; Matsushita et al. 2012), increase the complexity of genomes, and provides an essential theoretical basis for the successful evolution of existing plants. According to this study’s results, these are polyploid materials formed by chromosome polymerization of three species. In the BC1 generation, five sexual reproductive modes produced genetic materials with highly diverse ploidy (2n=35-84) and breeding application. 1) Parthenogenesis pathway (n) produced weaker, less viable plants, and hence were not recommended for follow-up research. 2) The sexual reproduction pathway (n + n) produced 63.33% of individuals with fewer wild chromosomes but higher translocations; thus, it is a preferential route for developing addition/substitution and introgression lines. 3) The polyspermy pathway (n+n+n) produced self-fertile progeny, thus fulfilling all criteria for developing new lines. 4) The apomictic pathway (2n) is a way to fix heterosis, shorten breeding years, and quickly fix excellent genes. 5) 2n+n pathway is one of the three major evolutionary drivers of ploidy increase (Ramsey and Schemske 1998); thus, it is a practical route for developing multiple new polyploid species. 2n and 2n+n pathways produced vegetatively propagated sterile perennial clones, providing practical evidence of instantaneous polyploidy speciation through strong reproductive isolation by sterility barriers (Comai 2005). This mode of speciation reduces effective migration rates, facilitates species divergence, provides chances to reproduce multiple times, and provides enough time for developing complete genetic isolation at later stages to neo-polyploids (Baack 2005; Rodriguez 1996). Stable fertile hybrid species can develop through recombinational speciation. According to this model, partially sterile individuals can give rise to specific novel genotypes through backcrossing or interbreeding, which could restore fertility and attain partial reproductive isolation from parents through chromosomal or genic sterility barriers (Dion-Côté et al. 2015; Grant 1958; Stebbins 1957). Theoretically, new fertile hybrid species can originate quickly but remain weakly isolated from parents at early stages. Strong reproductive isolation through chromosomal and genic recombination barriers requires an extended period (Abbott et al. 2010). In this study, originated allotetraploids were self-fertile and did not show remarkable fecundity up to the first seven observed generations, and preferably mate with tetraploid inbreds (∼80% seeds fertility) rather than diploid maize (1.03-7.17% seed fertility), which are minority cytotype exclusion and assortative mating mechanisms exhibited by neo-polyploids for their short-term early survival (Mable et al. 2011). The most concerning traits in the evolution from teosinte to corn are apical dominance and the evolution of compact cob in maize. Maize has paired mature spikelets with outer soft leaves coverings; however, only a single spikelet matures with outer harder glumes in teosinte. The cob of recently originated allotetraploids (NsAt) resembling maize, having paired spikelets but with fewer seed rows and reduced length. Cob morphological traits are quantitative in nature and are under the control of more than or just five genes or regions (Stitzer and Ross-Ibarra 2018); thus, different allele doses and combinations impact cob phenotypic diversity. Tillering/apical dominance is mainly controlled by a single gene (*tb1*) with major effects (Doebley et al. 1997). The chromosome containing branching allele may have lost, or chromosome rearrangements may have purged the *tb1* locus in these allotetraploids. Further study of this material can explore genetic mechanisms of compact ear and apical dominance behavior evolution in cultivated maize. **Figure S17** puts forward a workable route to synthesize different maize germplasm and improve tetraploid maize inbreds using maize wild relatives.

### Chromosome changes showed complementary characteristics of maintaining genomes at the full fold

One of the main problems in establishing new polyploids is the instability of genetic compositions and persistent karyotype changes. The current study discussed the formation of allopolyploid species between maize and tetraploid perennial teosinte, a milestone for studying allopolyploid diversity. Results showed that although the chromosomes of maize and Z. perennis had adjusted, the selection of chromosomes among generations tended to be around 40. Simultaneously, 45S and 5S rDNA loci tend to be in stable pairs, indicating that the chromosome and genome variation for evolution at the population level have complementary characteristics of maintaining its full integrity and fold, which can be helpful in establishing successful allotetraploid lines, and is in line with cytogenetic features of rDNA across land plants, showing a significant positive relationship of rDNA loci with ploidy (Garcia et al. 2017; Macháčková et al. 2021). This property can be used in allopolyploid maize genetic breeding and developing unique polyploid maize varieties. Although it has continuous karyotype changes and genome instability, its genetic material and chromosomes have significant recombination characteristics, differences in genetic material retention, and loss. Results show that different synthetic lines of these nascent allotetraploids have continuing stability of karyotypes, which paves the way for the homology of intermediate development processes and reproducibility. However, different genetic compositions exist within the sister lines and generations, laying a foundation for establishing polyploid diversity. Such karyotype instability has also been reported in previous auto and allotetraploid maize hybrids (Kato and Birchler 2006; Shaver 1964; Shaver 1962a). Chromosome number changes generate additional chromosome copies to enhance the genomic plasticity of polyploids (Stebbins 1971), which is a potential mechanism for fueling evolution. However, large-scale numerical chromosome instability with extensive rearrangements adversely affects fitness and may lead the polyploids to extinction. Only finely tuned individuals with balanced chromosomal variation can survive. The nascent near-allotetraploids in the current study had a narrow range of chromosomal number variations (38-42), indicating that more aberrant individuals (2n < 38 and >42) might be selected against. The chromosome range was further narrowed down in reconstructed tetraploid maize populations from F1 to F3 (Table 1), showing progressive genome stability. Interestingly, chromosomes numbers are preferably normalized to 2n=40. All *Zea* species have chromosomes multiples of x = 10; even maize has reduced down to n = 10 from an ancestral n = 20. The *Tripsacum* genus has maintained higher, albeit not a multiple of 10, numbers of chromosomes after the shared polyploid event.

### Uniparental chromosomes loss reflected genomic affinities between subgenomes

In nascent allotetraploids, almost all *Tripsacum* chromosomes were eliminated in early generations. *Tripsacum* is distantly related to maize, with only 1-4 chromosomes synteny with maize chromosomes, while maize and *Z. perennis* share the same genus (Galinat 1973; Iqbal et al. 2018; Longley 1941; Molina et al. 2006; Wet and Harlan 1974); thus, their chromosomes recombined, evolved, and sustained. Still, the detection of *Tripsacum* genetic transfer (by AFLP) and chromosome translocation in the current study revealed the possibility of genetic exchange between two genera, which is per previous studies (De Wet and Harlan 1974; Harlan and De Wet 1977; James 1979). Further partial preferential uniparental chromosome loss occurred for *Z. perennis* chromosomes when nascent near-allotetraploids hybridized with tetraploid maize inbreds (Table 5; Suppl. Fig S*12* – 16 and Suppl. Table S14). Complete uniparental chromosome elimination has also been previously reported in interspecific hybrids of closely related species (Finch 1983; Kasha and Kao 1970; Subrahmanyam 1977) and in the hybrids of distantly related species (Chen et al. 1991; Laurie and Bennett 1986; Matzk 1996; Rines and Dahleen 1990; Zenkteler and Nitzsche 1984). As observed for *Z. perennis*, partial uniparental chromosome elimination has also been reported in previously studied polyploids (Barclay 1975; Riera-Lizarazu et al. 1996). Several mechanisms of uniparental chromosome elimination reported in other polyploidy species can explain this phenomenon (Akera et al. 2017; Ishii et al. 2016; Ishii et al. 2010; Majtánová et al. 2021). The mechanism of uniparental/partial chromosome elimination in the current study requires detailed meiotic process elaborations for further understanding.

### Chromosome rearrangements in synthetic allotetraploid populations and establishment of polyploid lines

The structural chromosomal changes include rearrangements and genetic fragment exchange between homoeologous, chromosomes breakage, and rDNA loci changes, observed in synthetic (Pontes et al. 2004; Song et al. 1995; Xiong et al. 2011) and natural polyploid species (Lim et al. 2008). Our data from GISH and FISH demonstrated that massive genomic rearrangements had occurred in maize *Z. perennis* tetraploids. Extensive rearrangements are deleterious and may lead to genomic instability and extinction (Otto 2007). However, chromosome rearrangements produce novel phenotypes, which may facilitate niche exploitation. There is no evidence of highly rearranged genomes being selected during the establishment of polyploid species. Recently, a pairing regulator locus *PrZ* has been reported in maize x *Z. perennis* hybrids (Poggio and González 2018), which may control homoeologous pairing or selection culled the plant with aberrant phenotypes and low fertility in early generations and favored only those plants which have stable chromosome numbers and rDNA loci, better pairing control, and advantageous chromosomes rearrangements. The nascent allotetraploids were propagated and maintained for up to 7 generations by self-breeding, regardless of pollen viability and genome stability, which showed plants with highly unstable and aberrant phenotypes could not survive. Natural selection against extreme individuals was operating extensively in early generations through a decline in pollen viability and seed set. This phenomenon can be further elucidated by extending the age of nascent allotetraploid. The observation of the current study provides an experimental explanation of the origin and existence of segmental allopolyploids. In the absence of strong pairing control, the homoeologous chromosomes with higher genetic and structural similarity can pair and recombine as autopolyploids. Changes in chromosome number and translocations are the products of homoeologous exchanges, which may compromise fertility due to duplicated and deficient gametes; no doubt, it occurs, but homoeologous exchanges (duplication, deletion and translocations) produce vast karyotypes diversity and novel phenotypes that are important substrates for natural selection to establish nascent polyploids and fuel species evolution. Some of favoring HEs can fix in population, promotes karyotype divergence, and suppress homoeologous pairings by reducing synteny along chromosomes.

### Allotetraploidization of autotetraploid maize was possible by homoeologous exchanges followed by natural selection

The allotetraploidization of maize can be accomplished by restructuring a maize genome so that its chromosomes do not pair with normal maize genomes (Doyle 1979). Breeders tried different ways to overcome barriers to developing true-breeding tetraploid maize due to its benefits of large stature(Yao et al. 2011), vigorous growth, higher biomass, and improved nutrition (Atlin and Hunter 1984; Randolph 1935) compared to diploid maize. Genome restructuring by hybridization for preferential pairing is an effective strategy to circumvent the problems to some extent (Doyle 1986; Shaver 1963). In the current study, first, we systematically induced chromosome rearrangements, accumulated favoring rearrangements in independent lines through self-breeding and recurrent selection, and then transferred the differentiated chromosomes into autotetraploid maize for genome restructuring for preferential pairing. Meiotic irregularities contribute to genome instability, chromosome diversity, and seed fertility (Lim et al. 2008; Madlung et al. 2005; Szadkowski et al. 2010), which are directly linked to the establishment and survival of newly formed polyploids. This genome-constructed tetraploid maize showed far fewer IV pairing configurations (average 5.19 for BP and 5.36 for BT) and meiotic abnormalities (Fig.9; Table S1-14) compared to previously reconstructed allotetraploid maize (average IV; 8.10, 7.60 and 7.31; three populations) for improving preferential pairing (Doyle 1986), as well as from ordinary autotetraploid maize (Iqbal et al. 2018; Longley 1924; Shaver 1962b). These observations align with the notion that introducing exogenous genetic materials accelerates autotetraploid fitness (Adams and Wendel 2005). Different homoeologous exchanges (HEs) may contribute to cytological diploidization (Burns et al. 2021). The speciation and divergence between extant natural species have been accompanied by chromosome rearrangements such as in rice, maize, and sorghum (Murat et al. 2010), *Brassica* (Cheng et al. 2013), *Cucumis* (Yang et al. 2014) and *Arabidopsis* (Lysak et al. 2006). Reduction in recombination between diverged chromosomes is a product of favoring HEs fixation, which may explain the reduction of tetravalent formation in this population. The role of HEs in genome structuring for creating diversity, subsequent genome evolution, and speciation in segmental allopolyploids has been extensively reviewed (Mason and Wendel 2020). It has been reported in allotetraploid peanuts (Bertioli et al. 2019), allohexaploids of *Brassica (Gaebelein et al. 2019),* and synthetic rice polyploids (Zhang et al. 2019). Apparently, homoeologous exchanges combined with a compensation mechanism operated for evolving rearranged karyotypes without loss of gene information (dosage balance mechanism) in these newly synthesized segmental allotetraploids, which is consistent with the prediction of the gene balance hypothesis (Birchler and Veitia 2007). In summary; (i) this material can broaden the genetic basis of allotetraploids of maize and enhance its adaptability, (ii) allotetraploidization of maize has been accomplished by hybridizing nascent near-allotetraploids into segmental allotetraploids lines by a recurrent selection breeding system, and (iii) nascent near-allotetraploids of S1 to S6 can be utilized create several different allotetraploid maize germplasm for breeding research of maize polyploids.

## Acknowledgments

This work is supported by the National Natural Science Foundation of China (32272035), the Science and Technology Projects of Sichuan Province(2022NSFSC0167), the Forage Breeding Projects of Sichuan Province during the 14th Five-Year Plan Period (21ZDYF2201-3), the Science and Technology Projects of Sichuan Province(2021NZZJ0009), the Sichuan Corn Innovation Team of National Modern Agricultural Industry Technology System (sccxtd-2020-02), and the National Program on Key Basic Research Project of China (973 Program, 2014CB138705).

## Disclosure Declaration

All the authors have read and approved the manuscript, and there is no conflict of interest among the authors

